# Competition for light color between marine *Synechococcus* strains with fixed and variable pigmentation

**DOI:** 10.1101/2025.01.17.633661

**Authors:** Louison Dufour, Laurence Garczarek, Francesco Mattei, Bastian Gouriou, Julia Clairet, Morgane Ratin, David M. Kehoe, Jef Huisman, Jolanda M. H. Verspagen, Frédéric Partensky

**Affiliations:** Sorbonne Université, CNRS, UMR 7144 Adaptation and Diversity in the Marine Environment (AD2M), ECOMAP team, Station Biologique de Roscoff (SBR), 29680 Roscoff, France; Institute of Microbiology, CAS Centre Algatech, 37901 Třebon, Czech Republic; Sorbonne Université, CNRS, UMR 7093 Laboratoire d’Océanographie de Villefranche (LOV), Institut de la Mer de Villefranche (IMEV), 06230 Villefranche-sur-Mer, France; Indiana University, Department of Biology, Bloomington, IN 47405, USA; University of Amsterdam, Department of Freshwater and Marine Ecology (FAME), Institute for Biodiversity and Ecosystem Dynamics, NL-1018 Amsterdam, the Netherlands

**Author notes:** **Correspondence:** Corresponding Author: F. Partensky.

**Keywords:** Marine picocyanobacteria, *Synechococcus*, pigment type, chromatic acclimation, competition model

## Abstract

Competition between phytoplankton species for light has triggered an extensive diversification of photosynthetic pigments. In *Synechococcus* cyanobacteria, three major pigment types occur in the ocean: blue light (BL) specialists that have a high ratio of the BL-absorbing chromophore phycourobilin (PUB) to the green light (GL)-absorbing chromophore phycoerythrobilin (PEB), GL specialists that have a low PUB:PEB ratio, and cells that modify their PUB:PEB ratio to match the ambient color, a process called ‘Type IV chromatic acclimation’ (CA4). The abundance of CA4-capable cells in marine ecosystems suggests that CA4 confers a fitness advantage in certain light conditions compared to cells with fixed pigmentation. This hypothesis was tested by performing mono- and co-cultures of a BL specialist, a GL specialist and a CA4-capable strain in chemostats under different light conditions. Monocultures enabled us to parameterize a resource competition model that was used to predict competition between the three pigment types in co-cultures. In line with the model predictions, the BL specialist won in low blue light and the GL specialist won in low and high green light. Interestingly, we found that while the CA4-capable strain was at a disadvantage at low light, it was able to outcompete specialists in high blue light.

**IMPORTANCE:** *Synechococcus* cyanobacteria are ubiquitous and abundant in the lit layer of most marine ecosystems. This ubiquity relies in part on the wide pigment diversity of their light-harvesting complexes, with three main pigment types thriving in open ocean waters: green light specialists, blue light specialists and chromatic acclimaters, the latter being capable of matching their pigment content to the ambient spectral field. Here, we simulated the competition for light color that occurs between these pigment types in the field by co-culturing them in various light color and intensity conditions, and compared the resulting data to that of a competition model. This study provides new insights on how this key group of phytoplankton colonized the various spectral niches of the marine environment.

## INTRODUCTION

Although marine phytoplankton account for approximately half of global net primary production, their contribution is highly variable spatially depending on community composition, which is greatly influenced by local environmental conditions (1, 2). Despite their apparent continuity, the world’s oceans indeed encompass many different ecological niches delineated by temperature gradients and changes in the relative availability of two types of essential resources for which phytoplankton taxa compete: light and nutrients. Light, which provides the energy required for photosynthesis, fluctuates both quantitatively and qualitatively in the water column. The exponential decrease of light irradiance with depth is accompanied by a progressive narrowing of the visible light spectrum because of the absorption of different wavelengths by water, dissolved organic matter, particles and phytoplankton. The spectral field also strongly varies horizontally from coastal particle-rich to clear open ocean waters (3, 4). In this context, Holtrop *et al.* (2021) recently defined five distinct spectral niches in aquatic ecosystems (violet, blue, green, orange and red niches), with the last two being restricted to freshwater, estuaries and near-coastal environments. Competition for light occurring in these five niches has triggered a wide diversification of photoprotective pigments, which are needed to cope with strong irradiances in the upper euphotic layer, and the photosynthetic pigments used to collect Photosynthetically Active Radiation (PAR). This pigment variability allows different phytoplankton taxa to coexist by spectral niche differentiation, that is, by collecting distinct parts of the visible light spectrum (6, 7).

A striking example of this pigment diversification can be seen in marine *Synechococcus*. With an estimated global abundance of 7 × 10^26^ cells, this ubiquitous picocyanobacterium is the second-most abundant organism in the world’s oceans (8) and exhibits the largest pigment diversity within a single phytoplankton lineage known so far (9–11). Like most other cyanobacteria, *Synechococcus* uses large, hydrophilic light-harvesting complexes called phycobilisomes (PBSs) to collect photons and transfer their energy to photosystems (PS) I and II (12, 13). *Synechococcus* PBSs are comprised of a core from which extend six to eight peripheral rods (14, 15). Both core and rods are made of α-β heterodimers of phycobiliproteins held together by linker proteins. While the PBS core is predominantly composed of allophycocyanin (APC, maximum absorption wavelength *λ_max_* ≈ 650 nm), rods can be made of either phycocyanin (PC, *λ_max_* ≈ 620 nm), or a combination of PC and one or two additional types of phycoerythrin (PE-I and PE-II, *λ_max_* ≈ 560 nm; (9, 13). Each phycobiliprotein α-β dimer covalently binds between two (APC) and six (PE-II) chromophores at the level of highly conserved cysteine residues, an attachment generally catalyzed by phycobilin lyases (16). In marine *Synechococcus*, three distinct types of chromophores have been reported: phycocyanobilin (PCB, *λ_max_* ≈ 630 nm), phycoerythrobilin (PEB, *λ_max_* ≈ 550 nm) and phycourobilin (PUB, *λ_max_* ≈ 495 nm; (14)).

Three main pigment types (PTs) have been defined among *Synechococcus* strains based on the phycobiliprotein composition of PBS rods, with PT 1 possessing only PC, PT 2 containing both PC and PE-I, and PT 3 having PC, PE-I and PE-II. PT 3 was further divided into subtypes based on the relative cell content of PUB and PEB, as estimated by the relative ratio of fluorescence excitation at 495 and 550 nm (*Exc*_495:550_), with emission set at 580 nm. More precisely, the PUB:PEB ratio is low (*Exc*_495:550_ < 0.5) in subtype 3a, high (*Exc*_495:550_ ≥ 1.6) in subtype 3c and variable in subtype 3d (9, 10). The pigmentation of PT 3a and 3c strains remains fixed in changing light qualities, so they are often referred to as ‘green light (GL) specialists’ and ‘blue light (BL) specialists’, respectively (17). In contrast, PT 3d strains are capable of reversibly modifying their *Exc*_495:550_ ratio from 0.6-0.7 in GL to 1.6-1.7 in BL, a process called ‘Type IV chromatic acclimation’ (hereafter CA4; (18–20). Two genetically distinct types of CA4-capable strains (PTs 3dA and 3dB) have been described (10, 21), each possessing a small genomic island specifically involved in this process, and called the ‘CA4-A’ or ‘CA4-B’ island, respectively. Interestingly, it has been suggested that during evolution the acquisition of the CA4-A island conferred the ability to chromatically acclimate to GL specialists, and conversely that the CA4-B island provided the same ability to BL specialists (22–25).

The ecological significance of the CA4 process has long remained elusive due to the lack of an effective method to discriminate CA4-capable strains from BL and GL specialists. However, using three different PBS gene markers, Grébert and co-workers (26) were able to quantify the relative abundances of the different *Synechococcus* PTs throughout much of the world’s oceans by using metagenomic data from the *Tara* Oceans expedition. These authors demonstrated that CA4-capable cells accounted for ca. 41.5 % of the total marine *Synechococcus* population, with PT 3dA and 3dB cells being almost equally abundant (22.6 % and 18.9 %, respectively) but colonizing complementary ecological niches in the field. PT 3dA cells dominated in temperate and high latitude waters, while PT 3dB cells were more abundant in warm tropical waters. By comparison, the GL and BL specialists were found to represent 20.3 % and 33.4 % of the global *Synechococcus* population, respectively. The GL specialists were shown to be more abundant in near-coastal particle-rich waters where green light predominates, while the BL specialists prevailed in clear open ocean waters where blue wavelengths dominate the underwater light field (5, 26, 27). The prevalence of CA4-capable strains in marine ecosystems suggests that, in some light conditions, CA4 may confer a fitness advantage to cells that allows them to co-occur with, and even sometimes outcompete, cells with fixed pigmentation.

To test this hypothesis, we performed continuous mono- and co-cultures of a BL specialist, a GL specialist and a CA4-capable strain (belonging to PT 3dB), all coming from the same location in the Red Sea (Table 1). These experiments were performed in chemostats (28) under different conditions of light quantity (15 and 75 μmol photons m^−2^ s^−1^) and quality (BL and GL). Additionally, a series of photo-physiological measurements were carried out on all strains to determine their respective adaptive value and better understand the outcomes of competition experiments. Monocultures enabled us to parameterize a resource competition model, which was subsequently used to investigate competition for light between the three *Synechococcus* strains. The theoretical predictions were afterwards confirmed by conducting co-cultures with all three representatives. Altogether, we demonstrated that while the CA4-capable strain was at a disadvantage at low light, it was able to outcompete cells with fixed pigmentation in high blue light.

**Table 1.**
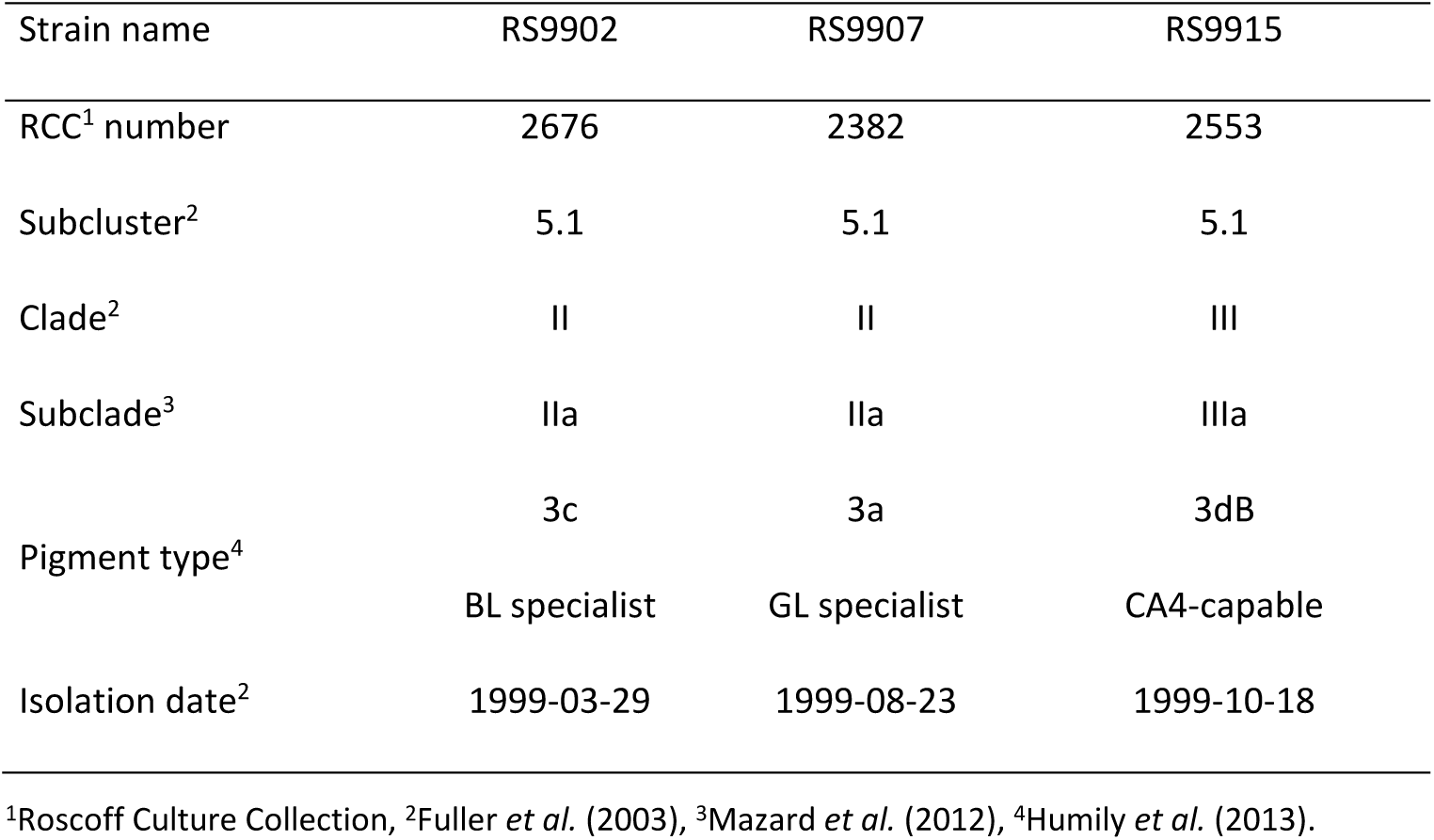
Characteristics of the three *Synechococcus* strains used in this study.

## COMPETITION MODEL

The resource competition model describes competition for light of different colors among phytoplankton species with different pigments. The model extends earlier competition models by Stomp and colleagues (6, 27) and was previously described (29). The model assumes a well-mixed water column that is illuminated from above with an incident light spectrum, *I_*i*n_*(*λ*), where *λ* represents wavelength. The underwater light spectrum *I_in_*(*λ*, *z*) changes qualitatively with water depth *z* due to absorption of different wavelengths by phytoplankton, water, dissolved organic matter (‘gilvin’) and suspended particles (‘tripton’). At each wavelength, light intensity diminishes exponentially with depth according to Lambert-Beer’s law, so that the underwater light spectrum over depth can be represented as:

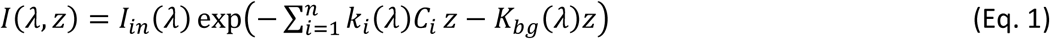

Here, *k*_*i*_(*λ*) represents the absorption spectrum of phytoplankton species *i*, and *C*_*i*_ is the population density of species *i*. The summation term signifies light absorption by *n* different phytoplankton species, each having its own distinct absorption spectrum. The absorption of photons by water, gilvin and tripton is incorporated in the wavelength-specific background turbidity *K*_*bg*_(*λ*). Furthermore, *I*_*out*_(*λ*) is defined as the light spectrum at the bottom of the water column, so that *I_out_*(*λ*) = *I*(*λ*, *z*_*m*_), where *z*_*m*_ is the water column depth.

The population density of each phytoplankton species *i* changes dynamically through growth and loss processes:

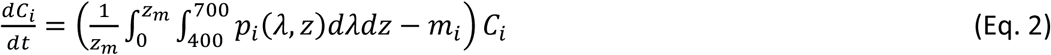

where we have *I* = 1, …, *n* different species in the community, *p*_*i*_(*λ*, *z*) is the specific production rate of species *i* as a function of wavelength *λ* and depth *z*, and *m*_*i*_ is the specific loss rate of species *i*. Since our experiments were performed at relatively low, non-saturating light intensities, we here assume that the specific production rate depends linearly on the quantity of photons absorbed by species *i* and the efficiency with which these photons are used:

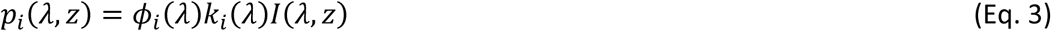

where *ϕ*_*i*_(*λ*) is the photosynthetic efficiency, i.e. the efficiency with which species *i* converts absorbed photons into production of biomass. The photosynthetic efficiency may vary with wavelength (29).

With some algebra, the depth integral in Eq. 2 can be solved with the help of Eq. 1 and Eq. 3. We then obtain:

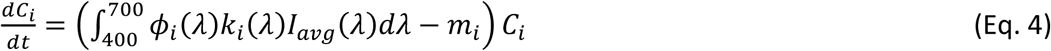

where the depth-averaged light intensity in the water column, *I_avg_*(*λ*), is defined as:

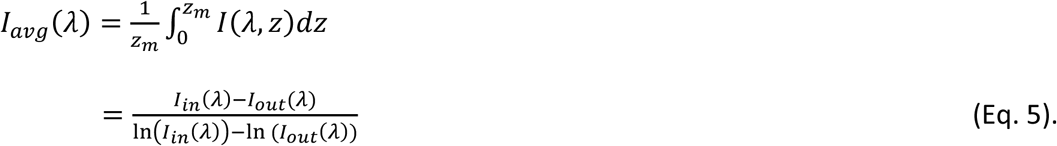

In our experiments, we used only two colors, BL and GL, rather than the full light spectrum. In this case, the population dynamics in Eq. 4 can be simplified to:

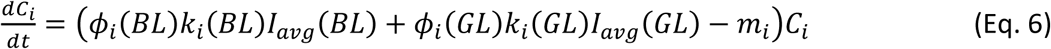

where the specific production rate is summed over the two colors BL and GL to obtain the total growth rate of species *i*.

According to this competition model, the species interact with each other by their modification of the underwater light spectrum. That is, if the population density of a species increases, then it diminishes the intensity and alters the spectral composition of light (via Eq. 1). These changes in the underwater light field, in turn, affect the population dynamics of itself and the other species in the community (via Eqs. 5 and 6).

## EXPERIMENTAL PROCEDURES

### Biological material and culture conditions

Three *Synechococcus* strains, a GL specialist (RS9907, PT 3a), a BL specialist (RS9902, PT 3c) and a CA4-capable strain (RS9915, PT 3dB; Table 1) were obtained from the Roscoff Culture Collection (https://roscoff-culture-collection.org/). All were isolated from the upper mixed layer at Station A in the Gulf of Aqaba (Red Sea) in 1999, but at different seasons (30). Cells were grown in PCR-S11 medium (31) supplemented with 175 µM K_2_HPO_4_,3H_2_0 and 2 mM NaNO_3_. Before starting the experiments, cultures were pre-acclimated for at least one month in temperature-controlled chambers at 25°C to the four different light conditions tested in this study: low blue light (LBL), low green light (LGL), high blue light (HBL) and high green light (HGL). Low (LL) and high light (HL) conditions corresponded respectively to 15 and 75 μmol photons m^−2^ s^−1^. Continuous light was provided by blue and/or green LEDs (Alpheus). Spectra of incident light and maximum emission wavelengths (*λ_max BL_* = 475 nm and *λ_max_* _*GL*_ = 515 nm) are provided in Figure S1. It should be noted that the green LEDs used in this study peaked in the valley between the PUB and PEB excitation peaks. Yet, in a separate study (21), we found that when acclimated to these green LEDs, all tested CA4-capable strains (including RS9915) exhibited the minimal *Exc*_495:550_ ratio expected for such CA4-capable cells, demonstrating that they sensed this light color as being ‘pure GL’.

For each light condition, continuous mono- and co-cultures of the three *Synechococcus* PTs were grown in chemostats (28) inoculated at an initial cell density of 3 × 10^6^ cells mL^-1^ of each strain, so that co-cultures started with an initial concentration of 9 × 10^6^ cells mL^-1^. Cells were grown in Pyrex® Roux flasks (SCI Labware) and continuously diluted until the culture reached the steady state, that is, at least five consecutive days with less than 10% variation in cell density (32). Each flask was equipped with a 5-inlet silicone cap for i) passive ventilation through a 0.2 µm air filter (Midisart 2000, Sartorius), ii) continuous supply (41.7 µL min^-1^, corresponding to a dilution rate of 0.1 day^-1^) of PCR-S11 medium using peristaltic pumps (Ismatec Reglo ICC, Cole-Parmer), iii) continuous removal of the overflow so that the volume in flasks remained constant over time, iv) aeration with 3% CO_2_-enriched air to control the pH of the cultures and avoid carbon deficiencies, and v) sample collection (Fig. 1). All experiments were done in biological duplicates, unless specified otherwise.

**Fig 1.**
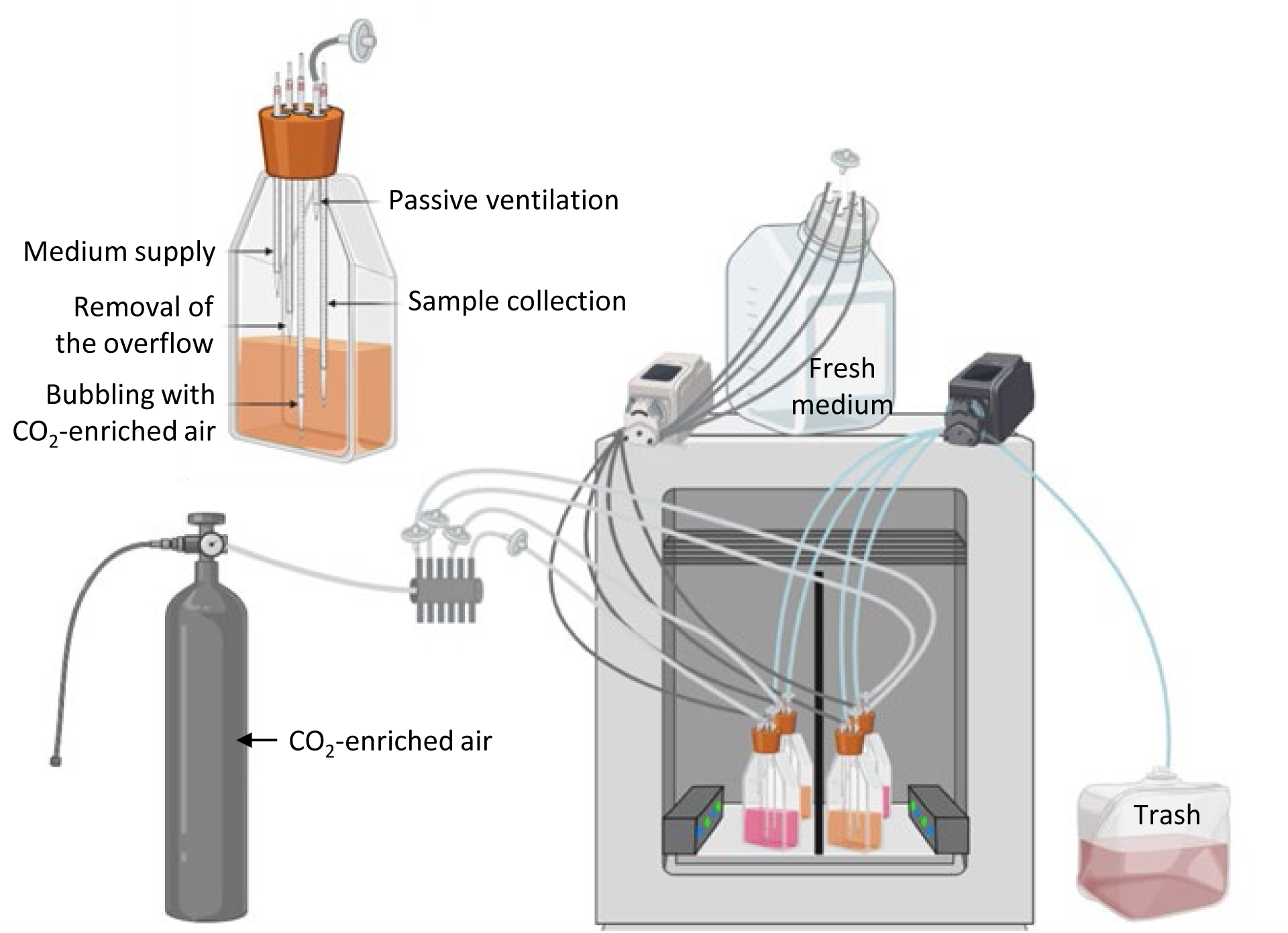
Experimental set-up for continuous monocultures and co-cultures in chemostats. The insert in top left corner is a zoom on one of the flasks shown in the general view.

For each mono- and co-culture, samples were harvested every one or two days to measure cell concentration, PSII quantum yield, chromophore and phycobiliprotein ratios, light intensity and pH. In addition, 5 mL aliquots from each of the co-cultures were collected at the same frequency, filtered through 0.2 µm Supor filters (Pall Life Sciences) and kept at −80 °C until analyzed by real-time quantitative PCR.

### Flow cytometry

Culture aliquots were sampled every day, fixed with 0.25% (v/v) glutaraldehyde (grade II, Sigma Aldrich) and stored at –80°C until analysis (33). *Synechococcus* cell densities were determined using a Guava easyCyte flow cytometer equipped with a 488 nm laser and the Guavasoft software (Luminex Corporation).

### PAR measurements

Before starting the experiments, the integrated intensity (over 400-700 nm) of the incident light (*I_in_*), as well as the light transmitted through the culture flasks (*I_out_*) filled with fresh PCR-S11 medium only, were measured with a PG200N Spectral PAR Meter (UPRtek) at 5 different locations within flasks, and the values averaged. During the experiments, *I_out_* was measured daily in the same way.

### Fluorimetry

#### Photosystem II quantum yield

Measurements of the PSII quantum yield (*F*_*V*_/*F*_*M*_), a proxy for the maximum photosynthetic activity of the cells, were carried out three times per week with a multi-wavelength fluorometer Phyto-PAM II (Walz), as previously described (34) except that five wavelengths (440, 480, 540, 590, or 625 nm) of modulated light were used.

#### PSII cross-section

The PSII cross-section (*σ*(*II*)_*λ*_), which represents the PSII functional antenna size (35), was measured twice during the experiments (growth phase and steady state) and only in monocultures. The O-I1 fluorescence fast kinetics, which refers to the increase of fluorescence yield induced by a strong actinic light, was recorded with a Phyto-PAM II (Walz), as described elsewhere (36). PSII cross-section values were calculated using the Phytowin 3 software (Walz; Schreiber *et al.*, 2012) at 480 nm (cyan) and 540 nm (green), the two wavelengths closest to PUB (*λ*_*max*_ ≈ 495 nm) and PEB (*λ*_*max*_ ≈ 550 nm) absorption peaks, respectively.

#### PSII electron transport

Like the PSII cross-section, the linear electron transport rate through PSII (*ETRII*) was measured twice (in growth phase and in steady state) in monocultures at 480 and 540 nm. As described by (36), the basal fluorescence *F*_0_ was recorded before samples were exposed to 13 steps of increasing light irradiance (90 seconds each). At the end of each step, the instantaneous (*F*_*o*_) and maximal (*F*_*M*_^′^) fluorescence levels were recorded, allowing the computing of PSII quantum yield under illuminated conditions 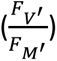. The *ETRII* at each step was calculated as follows:

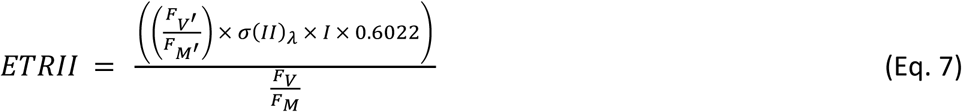

with *I*, instantaneous irradiance (37).

Finally, *ETRII* values for each step were plotted against their corresponding light irradiance. The photosynthesis Platt model (38) was then fitted to these curves and used to compute the PSII efficiency under non-saturating light (*⍺*; (36)).

### Spectrofluorimetry

*In vivo* fluorescence spectra were recorded several times a week using a FL6500 spectrofluorimeter (Perkin-Elmer) and analyzed with the Fluorescence software (Perkin-Elmer), as described elsewhere (21). Briefly, excitation spectra were acquired between 450 and 560 nm (with emission set at 580 nm) and emission spectra between 550 and 750 nm (with excitation set at 530 nm). The *Exc*_495:550_ fluorescence excitation ratio was used as a proxy of the whole cell PUB:PEB ratio. *Em*_560:650_ and *Em*_650:680_ fluorescence emission ratios were used to estimate the PE to PC as well as the PC to PBS terminal acceptor (TA) ratios, respectively. The PE:PC ratio provided insights into the electron transfer efficiency within the PBS and the length of PBS rods, and the PC:TA ratio into the coupling between the PBS and PSII reaction center chlorophylls.

### Real-time quantitative PCR

#### Design of the probes and optimization

Because it was not possible to differentiate all pigment types by flow cytometry under some conditions, particularly under BL where the CA4-capable strain and the BL specialist had indistinguishable fluorescence signals, a real-time quantitative PCR approach was developed to assess the relative abundance of each pigment type within the co-cultures. For each strain, primers were designed using Geneious® (version 11.0.5). Target genes were selected as being single copy and strain-specific, based on patterns of gene presence/absence in the Cyanorak information system (Table S1; (39)). Each set of primers was tested for specificity and optimized using DNA extracted from cultures of the three studied strains.

#### Sampling, cell lysis, DNA extraction and purification

Five mL of each co-culture were sampled daily, filtered through 0.2 µm Supor filters of 25 mm diameter (Pall Life Sciences) and stored in 2 mL Eppendorf tubes at –80°C until analysis. DNA was extracted from the 0.2 µm Supor filters following a protocol adapted from previous studies (40, 41). After thawing the filters on ice, 350 µL of lysis buffer (50 mM TRIS, 20 mM EDTA, pH 8.0) and a 5 mm steel bead were added to each Eppendorf. The samples were then grounded for 30 sec at 30 Hz using a TissueLyser (MM300, Retsch) to break the cell wall at room temperature. To ensure complete cell lysis, filters were incubated with 175 µL lysozyme (50 mg mL^-1^, Sigma-Aldrich) for 45 min at 37°C under agitation (Thermomixer comfort, Eppendorf). 70 µL of SDS (10%, Invitrogen) and 35 µL of proteinase K (20 mg mL^-1^, Sigma-Aldrich) were added and filters were again incubated under agitation for 2.5 h at 55°C. RNA was then removed by adding 70 µL of RNase (20 mg mL^-1^, Sigma-Aldrich) for 10 min at room temperature. The filters and the aqueous phases were immediately transferred to 2 mL Phase Lock GelTM (PLG) tubes (QuantaBio, VWR). To dissolve the filters, increase DNA extraction efficiency and remove protein contaminants, two phenol:chloroform:isoamyl alcohol (25:24:1 v/v; Eurobio) and one chloroform:isoamyl alcohol (24:1 v/v; Sigma-Aldrich) extractions were conducted. For each, 700 µL of organic extraction mix was added to the PLG tubes. After mixing, the tubes were subsequently centrifuged for 5 min at 1500 x *g* and 18°C using an Eppendorf 5417R. The aqueous phase was then recovered and transferred to new PLG tubes. Finally, the DNA in the aqueous phase was purified using silica gel columns (DNeasy Blood and Tissue Kit, Qiagen) following the manufacturer’s protocol for Gram-negative bacteria. DNA was eluted in 200 µL nuclease-free water (Invitrogen). 260:280 and 260:230 ratios were measured to assess the DNA purity using a ND-1000 spectrophotometer (Nanodrop).

#### Preparation of real time PCR standards from Synechococcus cultures

Template DNA was obtained from exponentially growing *Synechococcus* cultures of each strain harvested by centrifugation at 4°C, 14 000 x *g* for 10 min using an Eppendorf 5804R in presence of 0.01% Pluronic acid (Sigma-Aldrich). 200 µL of ATL buffer (Qiagen) was added to the pellets before grinding using a 5 mm steel bead, as described above. Then, samples were incubated under agitation (Thermomixer comfort, Eppendorf) with 10 µL lysozyme (50 mg mL^-1^, Sigma-Aldrich) for 45 min at 37°C. After addition of 200 µL of AL buffer (Qiagen) and 15 µL of proteinase K (20 mg mL^-1^, Sigma-Aldrich), samples were incubated under agitation for 2.5 h at 55°C. 200 µL ethanol 100% (Sigma-Aldrich) was immediately added to the samples, which were vortexed and purified using silica gel columns (DNeasy Blood and Tissue Kit, Qiagen) following the manufacturer’s instructions. DNA was eluted in 100 µL nuclease free water (Invitrogen). As for the co-culture samples, DNA 260:280 and 260:230 ratios and concentrations were quantified using a ND-1000 spectrophotometer (Nanodrop).

#### Real time PCR

Reactions for real time PCR were prepared by mixing 6 µL of ONEGreen® FAST qPCR Premix (Ozyme), 0.12 µL of forward and reverse primers stock (final concentration of 300 µmol L^-1^, Eurogentec), 0.76 µL of nuclease-free water (Invitrogen) and 5 µL of template DNA. Template DNA for the standard curves was serially diluted 10-fold over 7 orders of magnitude to obtain standard concentrations ranging from 50 to 5.10^-6^ ng per well. Template DNA from the co-culture samples was not diluted. Each reaction was performed in triplicate. Real-time SYBR Green fluorescence data were acquired using a LightCycler® 480 (Roche) and the program recommended by the manufacturer. The baseline and threshold cycle (C_t_) were estimated automatically with the ‘Abs Quant/2^nd^ Derivative Max’ of the LightCycler® 480 software (version 1.5.0, Roche). For co-culture samples, each quantification cycle (C_q_) was converted to DNA concentration based on the standard curve equation. The number of target gene copies per well was then estimated using Dhanasekaran *et al.*’s formula (42) and converted to copy per mL based on the volume of the filtered co-culture, the total amount of DNA extracted and the amount of template DNA added in each well. Results were expressed as percentage of each strain in the co-culture at any given time of the experiments.

### Comparative genomics

The repertoire of genes involved in the biosynthesis or regulation of PBS components was compared between the three *Synechococcus* strains using the Cyanorak genome database (www.sb-roscoff.fr/cyanorak/; (39)). The number of phycobilin lyases, the enzymes responsible for the covalent binding of chromophores to phycobiliproteins, encoded in each genome was then used to predict the chromophorylation of each α-β dimer based on literature on lyase function, while the PBS linker content was used to estimate the PBS rod length (15).

### Biovolume

For each tested light condition, the biovolume of each strain (Table S2) was estimated at steady state using an Eclipse 80i fluorescence microscope equipped with a Cy3 filter (Nikon). Photographs were taken at x100 magnification using a SPOT RT3 camera (Diagnostic Instruments Inc.) under GL excitation (*λ*_*max*_ = 550 nm) to observe their natural orange fluorescence. Analysis of the photographs was done with SPOT Advanced software (Diagnostic Instruments Inc.) to measure the length (*B*) and width (*W*) of 100-150 cells. The biovolume (*V*) was calculated assuming that *Synechococcus* cells have a short-rod shape, using the following formula:

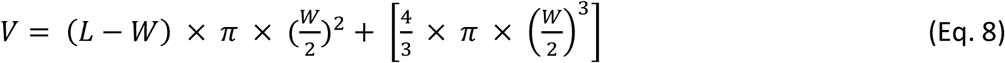

### Dissolved nutrient analyses

Phosphate (PO_4_^3-^), nitrate (NO_3_ ^-^) and ammonium (NH_4_^+^) concentrations were quantified in steady state following standardized protocols (43, 44).

### Statistics

Statistical analyses were carried out using R software (version 4.2.3; R Core Team, 2021) to compare the pigment and photosynthetic characteristics of i) a given strain in the different light conditions tested (LBL, LGL, HBL, HGL), and ii) the three different strains in a given light condition. One-way ANOVA or Kruskal-Wallis (stats package version 3.6.2) tests were performed, after checking for the normality (Shapiro Wilk’s test; stats package version 3.6.2) and variance homogeneity (Levene’s test; car package version 3.1-3) of the data, and followed by Tukey’s (stats package version 3.6.2) or Dunn’s tests (dunn.test package version 1.3.6), respectively. The same approach was used to compare critical light intensities, i.e. irradiances transmitted through the cultures at steady state (46, 47), for each PT representative and a specific light condition.

### Estimation of model parameters

All model parameters were estimated from monoculture experiments. System parameters such as the incident intensities of LBL, LGL, HBL and HGL (*I*_*in*,*lb*_, *II*_*in*,*lg*_, *II*_*in*,*hb*_, and *II*_*in*,*hg*_), the depth of the chemostat vessel *z*_*m*_, and the dilution rate *D* were defined by the experimental settings (Table 2). We assumed that the specific loss rates of the strains were determined by the dilution rate of the chemostat vessel (i.e., *m*_*i*_ = *D*). Light absorption coefficients (*k*_*i*_) of the three *Synechococcus* strains and background turbidity (*K*_*bg*_) were estimated for each combination of light color and light intensity (i.e., LBL, LGL, HBL and HGL). First, based on Eq. 1, for each monoculture we plotted the values of *ln*(*I_out,j_*/*I_in,j_*)/*z*_*max*_ versus *C*_*i*_ measured at each time point of the experiment. Linear regression analysis was then applied and the light absorption coefficient (*k*_*i*_) and background turbidity (*K*_*bg*_) were estimated as the slope and intercept of the linear regression. Finally, the estimates of *K*_*bg*_ and *k*_*i*_ obtained from these regression analyses were averaged over the replicates of each treatment. The photosynthetic efficiencies (*ϕ*_*i*_) of the three *Synechococcus* strains were estimated for each light condition by fitting the time courses of population density (*C*_*i*_) and light transmitted through the chemostats (*I_out_*) predicted by the model to the time courses observed in the monoculture experiments, using a least-squares method (47).

**Table 2.**
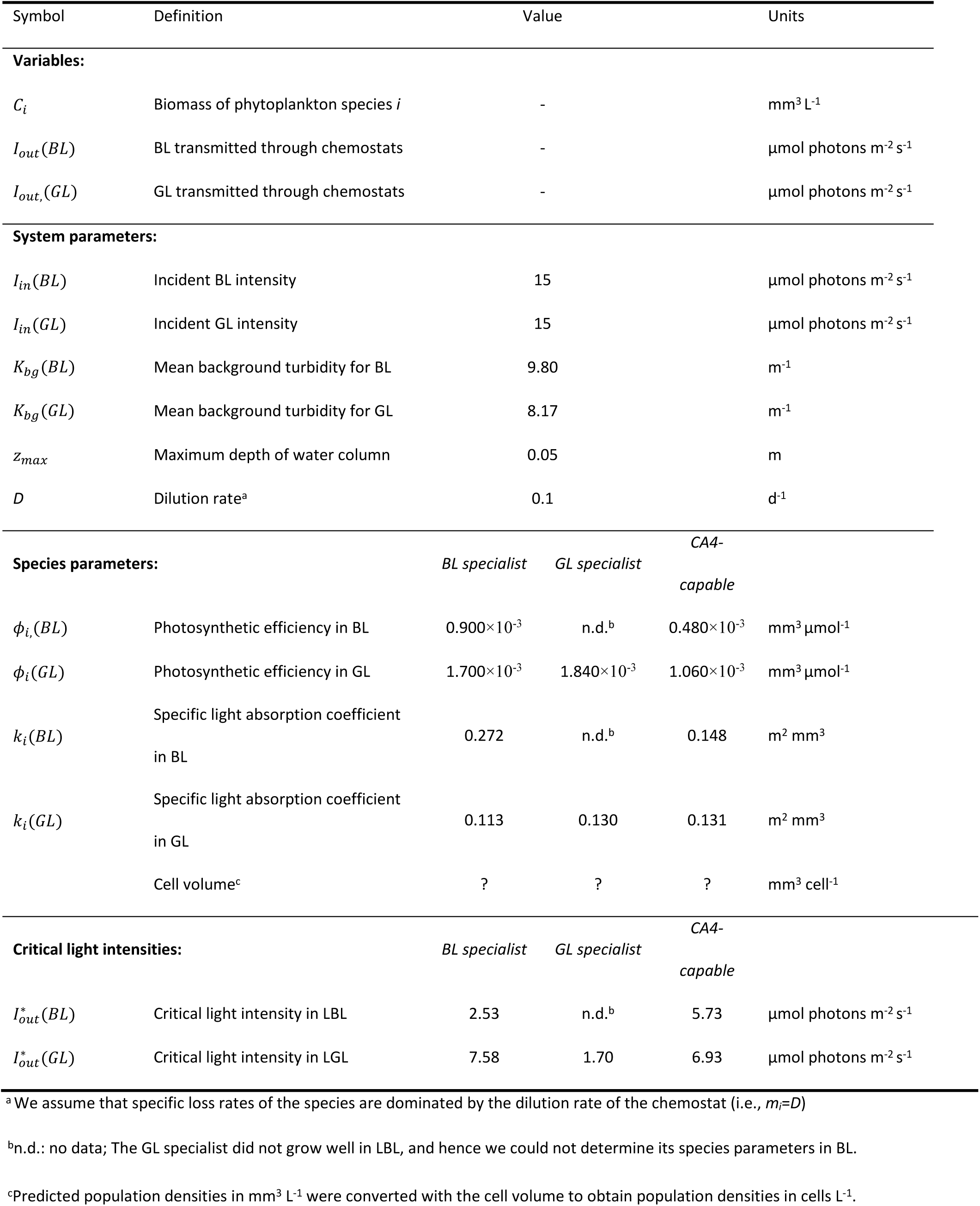
Model parameters estimated from the monoculture experiments. Abbreviations: BL, blue light; GL, green light; LBL, low blue light; LGL, low green light.

The parameter estimates obtained from the monoculture experiments are summarized in Table 2. These monoculture estimates were used to predict the time courses and outcomes of the competition experiments.

## RESULTS

### Monocultures

#### Time course of cell densities and light penetration in different light conditions

To better understand how each of the three main *Synechococcus* PTs thriving in the open ocean (26) individually behave in different conditions of light quantity and quality, we first performed continuous monocultures of one representative strain for each PT: RS9902, a BL specialist (PT 3c); RS9907, a GL specialist (PT 3a); and RS9915, a CA4-capable strain (PT 3dB; Table 1).

During the experiments, the cell density gradually increased to a steady state (Figs. 2A, 2D, 3A and 3D), causing a gradual decrease of the intensity of light transmitted through the culture flasks (*Iout*; Figs. 2B, 2E, 3B and 3E). Two parameters derived from these measurements, namely the cell concentration at steady state and the critical light intensity (*I*^\**out*^), i.e. the light intensity transmitted through the culture at steady state (46, 47), were used to fit the model. The PSII quantum yield (*F*_*V*_/*F*_*M*_), a proxy of the maximum photosynthetic activity of the cells, was also measured on a regular basis during the experiment in order to gain insights about the physiological status of the cells (Figs. 2C, 2F, 3C and 3F).

**Fig 2.**
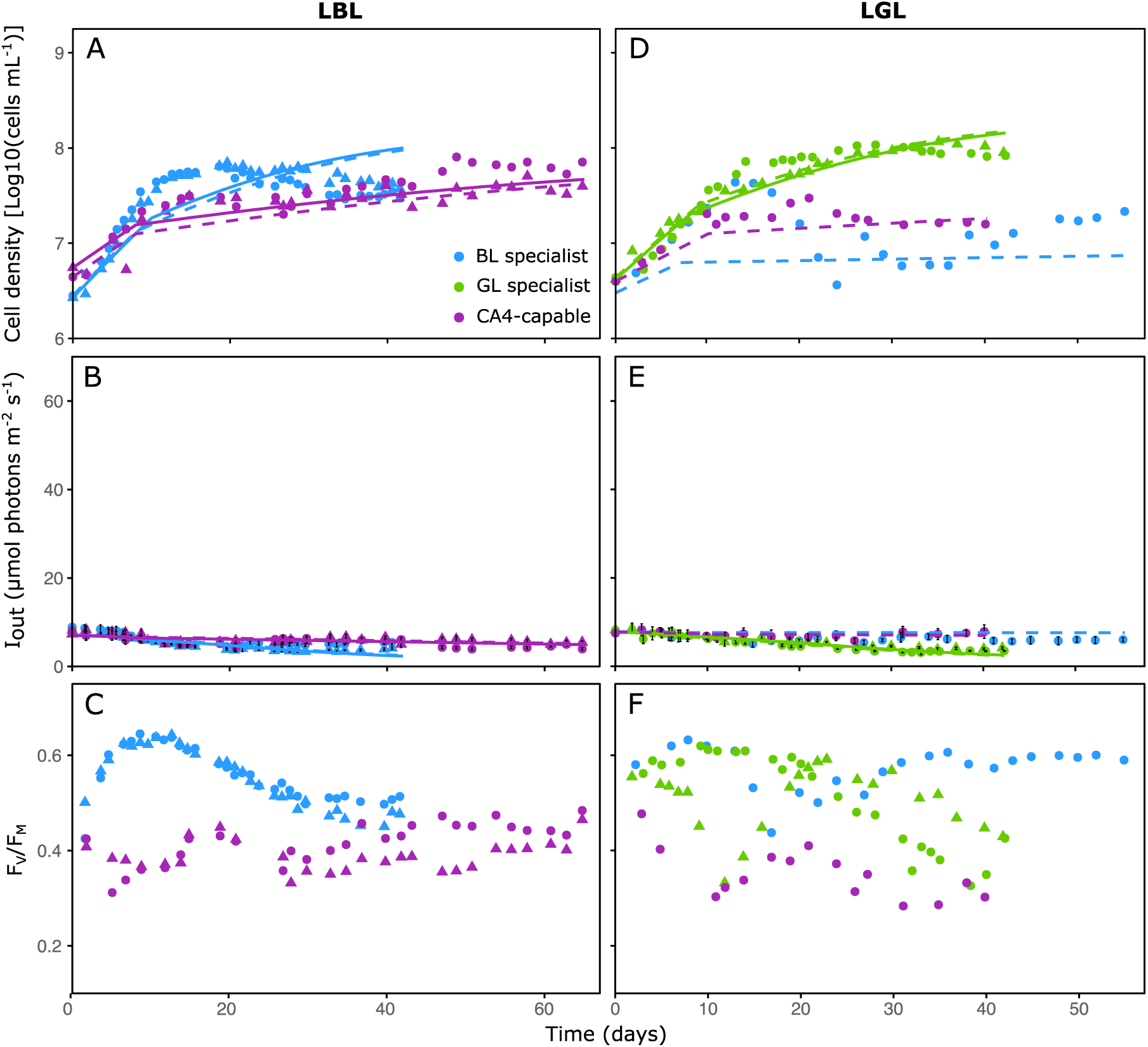
Monocultures of three marine *Synechococcus* strains representative of different pigment types in low blue and green light conditions. (A through C) Low blue light (LBL). (D through F) Low green light (LGL). (A and D) Cell density. (B and E) Light transmitted through the culture flasks (*I_out_*). (C and F) Photosystem II quantum yield (*F*_*V*_/*F*_*M*_). Error bars in panels B and E correspond to the average and standard deviation of five different measurements on the side of the flask opposite to the light source. Shapes indicate different replicates (circles for replicate A and triangles for replicate B). Dashed (for replicate A) and solid (for replicate B) lines represent the input of the model.

**Fig 3.**
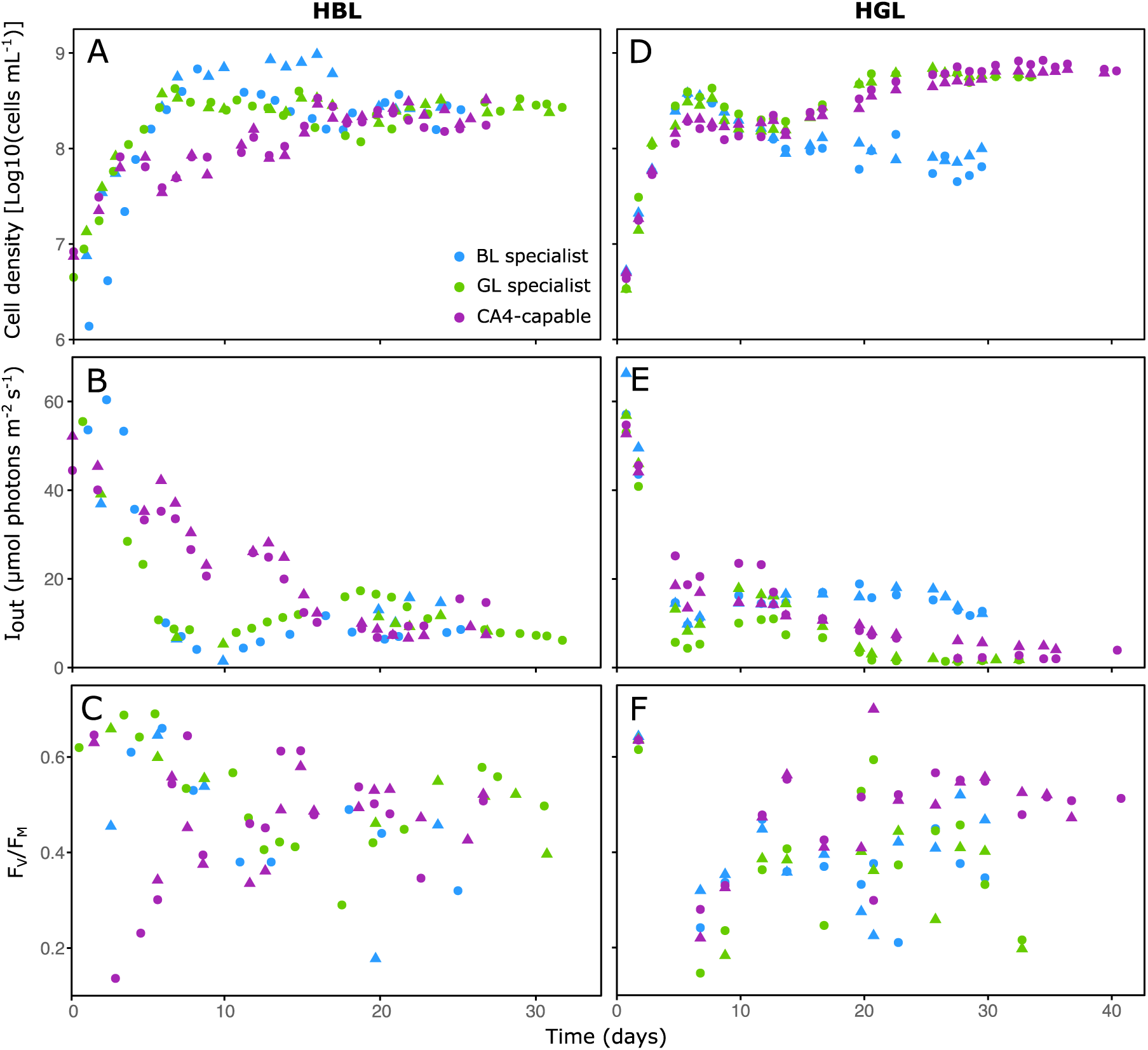
Same as Fig. 2 but in high blue light (HBL) and high green light (HGL) conditions.

In LBL, the BL specialist grew faster than the CA4-capable strain and reached steady state with cell densities of ≈3.6 × 10^7^ cells mL^-1^ by approximately the 35^th^ day of monitoring (Fig. 2A). In comparison, the CA4-capable strain stabilized more than 12 days later at similar or slightly higher cell concentrations (≈3.9 × 10^7^ cells mL^-1^ for one replicate and ≈6.4 × 10^7^ cells mL^-1^ for the other). Despite several attempts (n=5), the GL specialist was not able to grow in LBL, likely because its growth rate in this light condition was lower than the dilution rate. The *I*^*^_*out*_ was statistically similar for the BL specialist and the CA4-capable strain (4.46 ± 0.99 and 4.85 ± 0.85 µmol photons m^-2^ s^-1^, respectively; One-way ANOVA, p-value >0.05; Fig. 2B and Table S3). However, the CA4-capable strain exhibited a lower *F*_*V*_/*F*_*M*_ than its counterpart throughout the experiments (Fig. 2C).

In contrast to LBL, the GL specialist grew well in LGL and reached high cell concentrations in steady state (≈1.2 × 10^8^ cells mL^-1^; Fig. 2D). Conversely, the growth of the two other strains was clearly limited by light. In our single successful attempt (out of 5) to grow them in this light condition, the BL specialist and CA4-capable strain stabilized at much lower cell densities (≈1.9 × 10^7^ and ≈1.6 × 10^7^ cells mL^-1^, respectively) than the GL specialist. Consistently, the GL specialist also reached the lowest *I*^*^_*out*_ (3.70 ± 0.47 µmol photons m^-2^ s^-1^, as compared to 5.92 ± 0.12 and 7.61 ± 0.26 µmol photons m^-2^ s^-1^ for the BL specialist and CA4-capable strain, respectively; one-way ANOVA, p-value < 0.05; Fig. 2E and Table S3). As in LBL, the CA4-capable strain exhibited the lowest *F*_*V*_/*F*_*M*_ of all strains in LGL (Fig. 2F), suggesting that its photosynthesis was less efficient in LL compared to both specialists.

In both LBL and LGL, the model was in most cases able to correctly capture the dynamics of population density and *I_out_*, and in particular for the CA4-capable strain.

In HBL, the BL and GL specialists grew faster than the CA4-capable strain, both reaching their highest cell concentrations (≈9.6 × 10^8^ and 4.3 × 10^8^ cells mL^-1^, respectively) during the first two weeks of the experiments (Fig. 3A). However, the three PTs stabilized at similar cellular densities in steady state (≈2.2 × 10^8^ cells mL^-1^) and had statistically similar *I*^*^_*out*_ (10.43 ± 4.17 for the BL specialist, 8.60 ± 1.65 for the GL specialist, and 10.08 ± 3.04 µmol photons m^-2^ s^-1^ for the CA4-capable strain; Kruskal-Wallis, p-value >0.05; Fig. 3B and Table S3).

In HGL, although all three PTs grew at similar rates at the beginning of the experiments, only the GL specialist and the CA4-capable strain achieved maximum cell concentrations of ≈6.3 × 10^8^ cells mL^-1^ at steady state (Fig. 3D). In this light condition, the GL specialist reached the lowest *I*^*^_*out*_ (1.66 ± 0.22 µmol photons m^-2^ s^-1^, compared to 14.76 ± 2.25 and 3.84 ± 1.73 µmol photons m^-2^ s^-1^ for the BL specialist and the CA4-capable strain, respectively; Kruskal-Wallis, p-value <0.05; Fig. 3E and Table S3). In contrast to LL, all strains exhibited a strong variability of *F*_*V*_/*F*_*M*_ over the course of the experiments in HL, without any clear pattern (Figs. 3C and 3F).

In HL conditions, the model was only able either to capture the initial increase in population density but with an overestimation of the steady state population density (Fig. S2A), or to capture the steady state population density but with an underestimation of the initial increase in population density (Fig. S2B). These poor fits of the population dynamics caused similarly poor fits of the dynamics in *I_out_*, and this applied to almost all strains in both HBL and HGL. Hence, the model could not adequately describe the HL experiments in monoculture, and therefore was not applied to the HL competition experiments in co-culture.

#### Comparison of pigment and photosynthetic characteristics between the different strains

To better characterize the pigment and photosynthetic properties of each strain, and therefore better explain their distinct growth behaviors, additional measurements were performed on monocultures in the different tested light conditions.

The ratio of whole cell fluorescence excitation at 495 and 550 nm (*Exc*_495:550_), a proxy of the PUB:PEB ratio of the cells, remained constant during the experiments and did not differ between light conditions for both the BL specialist (1.58 ± 0.04) and the GL specialist (0.39 ± 0.01), as expected from their fixed pigmentation (Fig. 4A). Also as expected from Humily *et al.* (2013), the CA4-capable strain displayed a low *Exc*_495:550_ ratio in HGL (0.72 ± 0.07) and high *Exc*_495:550_ ratios in LBL (1.69 ± 0.06) and HBL (1.52 ± 0.07). Surprisingly, however, the *Exc*_495:550_ ratio of the CA4-capable strain varied during the time course of LGL monocultures, from a minimal value of 0.7 at the beginning of the experiment to an intermediate value of 1.03 ± 0.11 at steady state (Figs. 4A and S3). This suggests that the CA4-capable strain perceived a change in the light spectrum as the monocultures became denser, and accordingly adjusted its PUB:PEB ratio to optimize energy collection.

**Fig 4.**
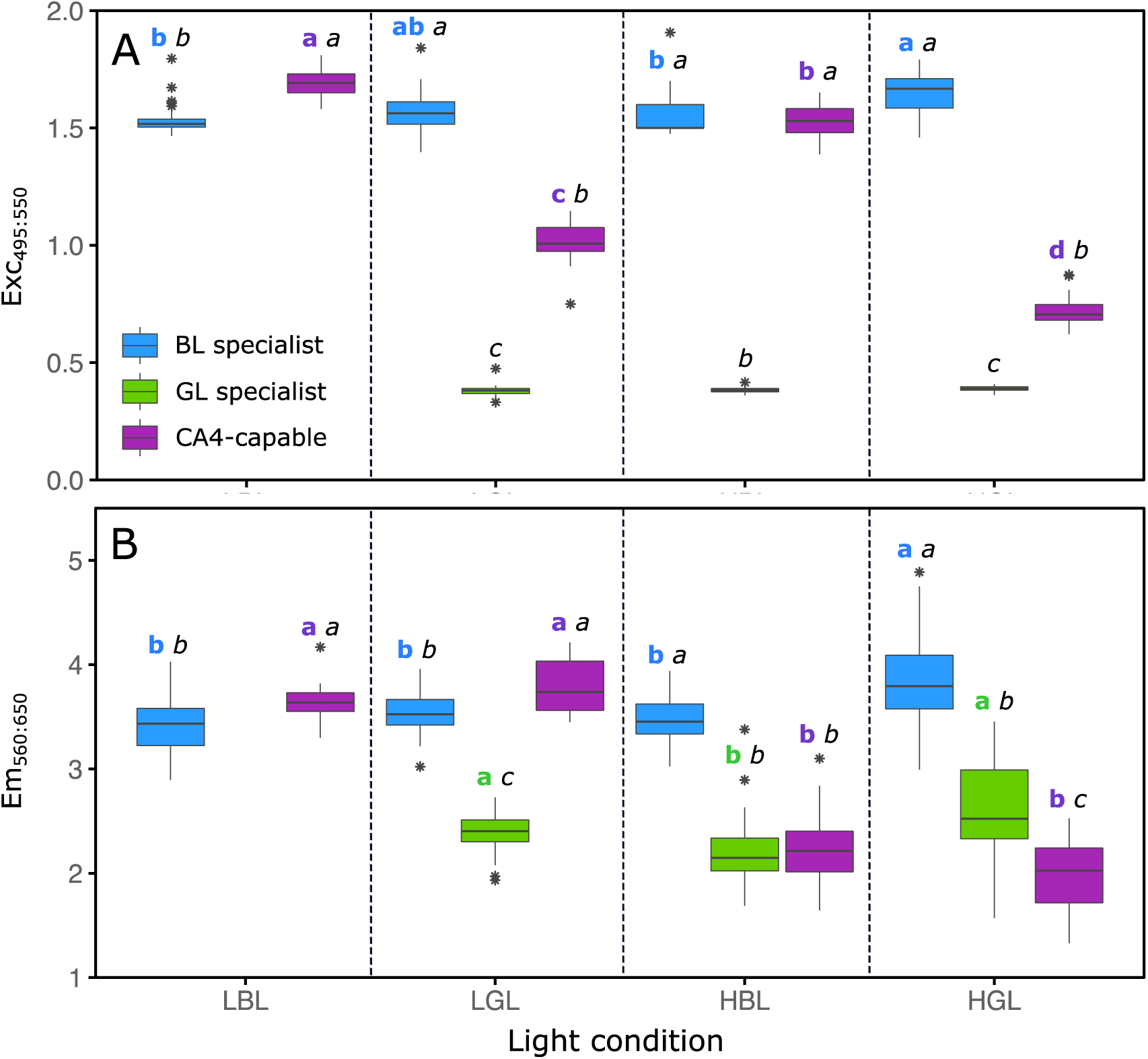
Mean *Exc*_495:550_ fluorescence excitation ratio and *Em*_560:650_ fluorescence emission ratio for the three *Synechococcus* strains grown in monocultures in the different light conditions used in this study. (A) *Exc*_495:550_ fluorescence excitation ratio, a proxy for the whole cell PUB:PEB ratio. (B) *Em*_560:650_ fluorescence emission ratio, a proxy for the whole cell PE:PC ratio. Boxplots represent all measurements performed over the course of the experiments. The different letters above boxplots indicate statistical test results (one-way ANOVA followed by Tukey’s test, or Kruskal-Wallis followed by Dunn’s test). The first bold colored letter compares the values displayed by a given strain in the four different light conditions tested. The second italic black letter compares the values exhibited by the three different strains in a given light condition.

To better interpret observed differences in PUB:PEB ratios between the strains, comparative analyses of their phycobilisome gene content, and more particularly of their phycobilin lyase gene content (Table S4), were made to predict the chromophorylation of phycobiliproteins at each cysteine binding site (Table S5), and thus to assess the molar PUB:PEB ratio per phycobilisome in both BL and GL (Table S6). These analyses predicted that the BL specialist and the BL-acclimated CA4-capable strain should exhibit the same molar PUB:PEB ratio (1.18), in agreement with their similar *Exc*_495:550_ ratio in LBL and HBL (Figs. 4A and S1). This implies that the different growth behaviors of the BL specialist and the CA4-capable strain recorded in LBL and HBL (Figs. 2A and 3A) cannot be explained by a difference in PUB:PEB ratio. In contrast, the molar PUB:PEB ratio of the GL specialist (0.22) is predicted to be almost twice lower than that of the CA4-capable strain (0.39) in GL, in agreement with the observed difference in *Exc*_495:550_ ratio between these strains (Figs. 4A and S1).

The ratio of whole cell fluorescence emission at 560 and 650 nm (*Em*_560:650_), a proxy for the phycoerythrin to phycocyanin (PE:PC) ratio of the cells, was systematically higher for the BL specialist than for the GL specialist, even though both exhibited some slight but statistically significant variations of their ratios depending on light conditions (Fig. 4B). Interestingly, the CA4-capable strain exhibited *Em*_560:650_ ratios similar to the BL specialist in both LL conditions, and to the GL specialist in HL conditions. By comparison, the ratio of whole cell fluorescence emission at 650 nm and 680 nm (*Em*_650:680_), a proxy of the PC to terminal acceptor (PC:TA) ratio of the cells, varied little between light treatments for all three strains although it was generally lower for the GL specialist than for the two other strains (Fig. S4).

As expected from the respective color preference of each PT, the PSII cross-section (*σ*(*II*)_*λ*_) as well as the PSII efficiency under non-saturating light (*⍺*) were both generally higher at 480 nm (cyan) than 540 nm (green) for the BL specialist, and conversely for the GL specialist (Figs. S5 and S6). The GL specialist was characterized by the highest *σ* and *⍺* values of all three strains at 540 nm in all light conditions where this strain grew, while the BL specialist exhibited the highest *σ* and *⍺* values at 480 nm in high light conditions only (Figs. S5 and S6). The CA4-capable strain displayed either low or intermediate *σ* values at both wavelengths, suggesting that its antenna size was smaller than specialists in their preferred color (Fig. S5). In addition, it generally had *⍺* values similar to (or even smaller than) specialists in their non-preferred color, suggesting that it had a fairly weak PSII efficiency under non-saturating irradiances (Fig. S6). A notable exception is the LBL condition, in which the CA4-capable strain exhibited higher *σ* and *⍺* values than the BL specialist at 480 nm.

### Competition experiments

Competition theory predicts that, if species compete for a single color of light, then the species with the lowest critical light intensity (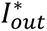) for this color will be the superior competitor (29, 46, 47). Hence, the 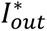 values estimated from the monoculture experiments can be used to predict the winners of a competition for light. The 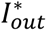 of the LL monocultures were estimated in two different ways: directly from the measurements of *I_out_* in monoculture at steady state (Table S3), and indirectly from the steady-state value of *I_out_* predicted by the model using the parameter estimates obtained from the LL monocultures (Table 2). Since the model could not adequately capture the population dynamics of the monocultures in HL, the 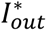 in these conditions were estimated only from the measurements of *I_out_* in monocultures. This resulted in the following predictions:

1. In LBL, the measured 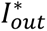 of the BL specialist and the CA4-capable strain were within the same range (Table S3), but the model analysis indicated that the 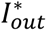 was lower for the BL specialist than for the CA4-capable strain (Table 2). Hence, the model predicted that the BL specialist should win.
2. In LGL, the measurements and the model analysis both showed that the 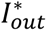 was lowest for the GL specialist (Tables 2 and S3). Hence, the GL specialist was predicted to win.
3. In HBL, the measured 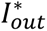 did not differ significantly between the three strains (Table S3), and hence were too close to reliably predict which of these three strains should be the best competitor in this light condition.
4. In HGL, the GL specialist had the lowest measured 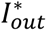 (Table S3), and hence was predicted to win the competition.

To test these predictions, continuous co-cultures were established with the three pigment types representatives in the same light conditions as for the monocultures. In LL, each of the BL and GL specialists won the competition in its favorite light color (Figs. 5A and 5D), while the abundance of the other PTs decreased. Therefore, the model was able to accurately predict the population dynamics during the competition experiments in both LL treatments (Figs. 5A and 5B). Although the temporal dynamics of the different strains in co-cultures was more variable between the two biological replicates in HL than LL treatments, the prediction proved valid for the HGL condition, where the BL specialist and the CA4-capable strain were completely outcompeted by the GL specialist at steady state (Fig. 5D). In HBL, where the model was unable to predict the outcome of competition, the co-culture experiment interestingly showed that the CA4-capable strain represented 85% to 90% of the total *Synechococcus* population at the end of the monitoring (Fig. 5C), the remainder being shared by the two specialists.

**Fig 5.**
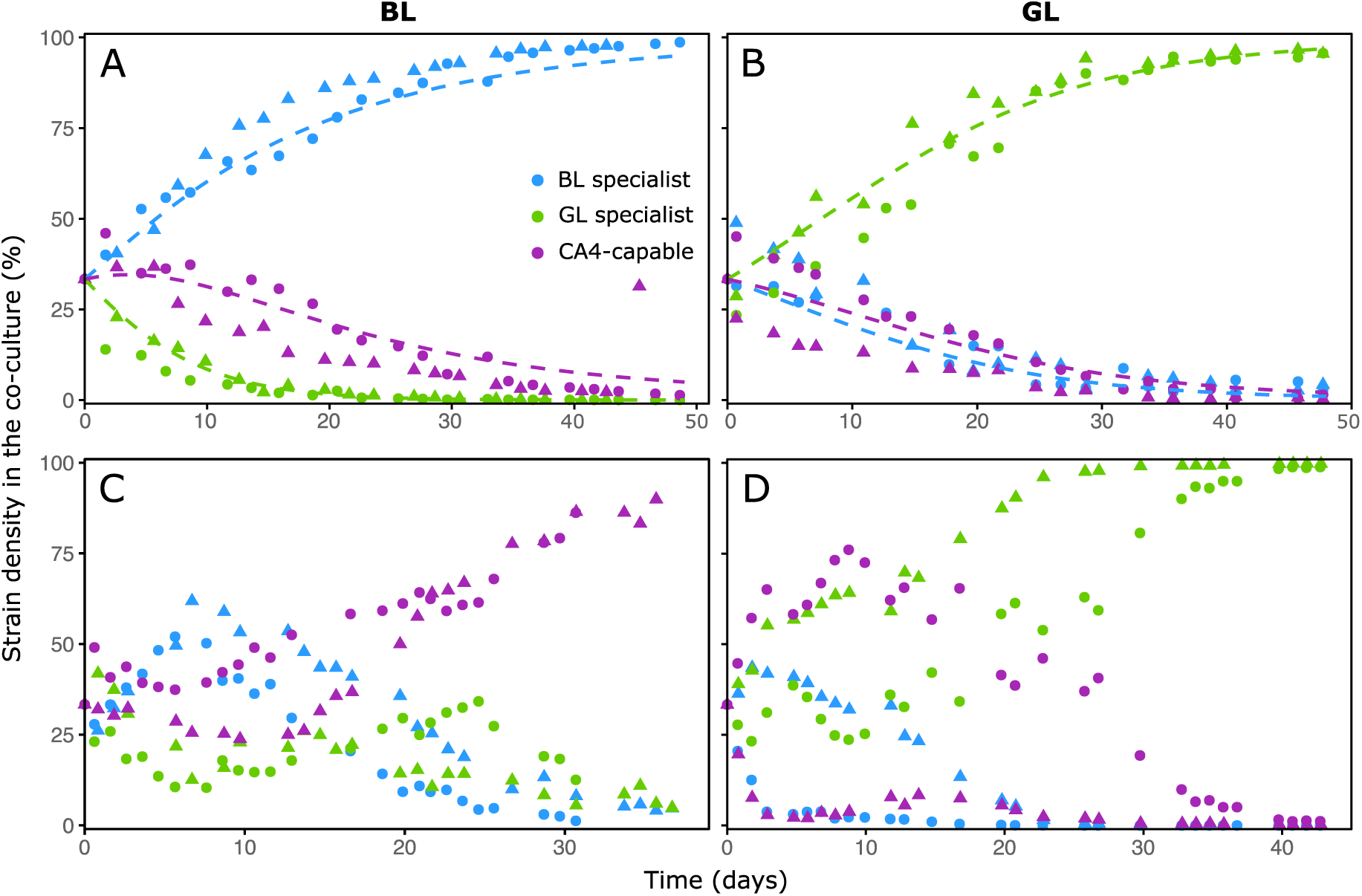
Time course changes of cell density in co-cultures of three marine *Synechococcus* strains representative of different pigment types in the four tested light conditions. (A) Low blue light (LBL). (B) Low green light (LGL). (C) high blue light (HBL). (D) High green light (HGL). Shapes indicate different replicates (circles for replicate A and triangles for replicate B). Dashed lines in panels A and B represent the input of the model.

### Nutrients

Quantification of NH_4_^+^, NO_3_^-^ and PO_4_^3-^ in steady state demonstrated that none of the mono- and co-cultures were nutrient-limited in LL (Table S7). By comparison, very sharp drops of NH_4_^+^ concentration were observed in HL cultures, with a majority of measurements below 50 μmol L^-1^ and a minimum value of 13 μmol L^-1^. As a consequence, all three strains assimilated nitrogen in the form of NO_3_ ^-^ (48), as indicated by the decrease in the NO ^-^ stock over the course of the experiments. However, NO_3_^-^ was sufficiently abundant at steady state in all cultures (above 900 μmol L^-1^) for the strains not to be nitrogen-limited. The concentration of PO_4_^3-^ also decreased in HL over the course of the experiments (minimum: 20 μmol L^-1^). However, it has been reported that marine *Synechococcus* strains grown in chemostats are able to grow at near maximum rates at PO_4_^3-^ concentrations below 10 nM (49). The conditions used by these authors being quite similar to ours (continuous culture, 24°C, 35 μmol photons m^-2^ s^-1^, 12h light-dark cycles), it is unlikely that our cultures faced phosphorus deficiency. These nutrient measurements demonstrate that, like for LL cultures, HL cultures were neither nitrogen-nor phosphorus-limited.

## DISCUSSION

While CA4-capable cells globally constitute the major *Synechococcus* PT in wide expanses of the world Ocean (26), the reasons for their ecological success are as yet unclear. One of the main current hypotheses, first evoked to explain the fitness advantage of a cyanobacterium exhibiting CA3 over either a red light specialist (PT 1) or a green-yellow light specialist (PT 2; (6, 50)), is that cells capable of chromatic acclimation would be better suited than specialists to sustain growth in a fluctuating underwater light field (51). Using a mathematical model of the ocean column, authors of the latter study found that deeper mixed layers selected for CA4-capable strains in simulated mixtures of *Synechococcus* PTs. However, results from their model did to match the actual distribution of PTs in the ocean, since they found no correlation between the relative abundance of CA4-capable strains and mixed layer depths, consistent with previous work (26). Alternatively, CA4-capable strains could perform better than specialists in certain light conditions. To test this hypothesis, we i) performed continuous monocultures of a BL specialist, a GL specialist and a CA4-capable strain in different conditions of light quantity and quality; ii) predicted the outcomes of competition experiments for light with a resource competition model; and iii) verified the theoretical predictions by co-culturing strains representative of all three PTs.

Results of co-culture experiments in LGL and HGL were consistent with competition theory, which predicted that the strain with the lowest 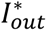 should be the superior competitor and thus displace its counterparts (46, 47). In both light conditions, the GL specialist displayed the lowest 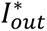 (Figs. 2E and 3E; Table S3) and was the only remaining strain at steady state in the co-cultures (Figs. 5B and 5D). This result was expected since the GL specialist was by far the best suited of the three PTs to harvest green photons. The GL specialist indeed not only possesses PBSs with a much lower molar PUB:PEB ratio (0.22) than the other two strains (0.39 for the CA4-capable strain in GL, and 1.18 for the BL specialist; Table S6), but also had by far the largest photosynthetic antenna size (Fig. S5B) and PSII efficiency under non-saturating light (Fig. S6B) of all three PTs at 540 nm.

The outcome of co-culture experiments in LBL also matched the model predictions since the BL specialist outcompeted the CA4-capable strain. This result might seem surprising since the latter strain exhibited a significantly larger antenna size (Fig. S5A) and PSII efficiency under non-saturating light (Fig. S6A) at 480 nm than the BL specialist. Yet, the CA4-capable strain also displayed a much lower PSII quantum yield (*F*_*V*_/*F*_*M*_) than its counterpart all over the course of the experiment (Fig. 2C), indicating that its photo-physiological status was suboptimal in LBL. This may partly explain why the CA4-capable strain was almost completely excluded by the BL specialist in the LBL co-culture (Fig. 5A). Regarding our inability to grow the GL specialist in LBL, our hypothesis is that the ambient photon flux in this light condition was too low to sufficiently feed its photosynthetic light reactions given the limited overlap between the wavelength range emitted by the blue LEDs and the fluorescence excitation spectrum of the GL specialist (Fig. S1A). This caused a rapid population loss due to continuous dilution (*D* = 0.1 day^-1^) that likely exceeded its growth rate. Such a scenario was previously evoked to describe the competition for light between two microalgae (28).

The outcome of co-culture experiments in HBL was more complex to interpret since the BL specialist and CA4-capable strain displayed very similar 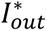 (Table S3), making it difficult to predict the best competitor based on these parameters. Photo-physiological measurements of monocultures revealed some interesting differences between the two PTs. On the one hand, the larger photosynthetic antenna size (Fig. S5A) and PSII efficiency under non-saturating light (Fig. S6A) of the BL specialist could *a priori* have conferred it a competitive advantage over the CA4-capable strain in HBL. On the other hand, the almost two times higher *Em*_560:650_ ratio of the BL specialist compared to the CA4-capable strain in this light condition (Fig. 4B) might reflect a lower energy transfer efficiency within PBS rods for the former strain, and thus explains the lower competitiveness of the BL specialist observed in this light condition. Given this uncertainty, the possibility that other factors are involved in the observed superiority of the CA4-capable strain in HBL cannot be excluded, though allelopathy can likely be excluded since we checked for the absence of microcin-C biosynthesis genes in both genomes (52). It is also worth noting that the high residual nitrate and phosphate concentrations (above 900 and 20 μmol L^-1^, respectively) indicate that nitrogen and phosphorus did not limit growth of the HL cultures. However, it is possible that the availability of some other elements (e.g., trace metals) became limiting or co-limiting in the very dense HL cultures, which may have affected competitive interactions between the strains. Future studies using different sets of BL-specialists and CA4-capable strains under a range of experimental conditions would also be needed to further test this finding elucidate in further detail which conditions favor CA4-capable strains, as well as to check whether a PT 3dA strain in competition experiments with specialists would behave similarly as the PT 3dB strain used in the present study, given their different evolutionary histories (24, 25).

Our results may have important implications for predicting the spectral niches of *Synechococcus* PTs in the world Ocean (5). Indeed, they imply that GL specialists should predominate in greenish environments, independent of depth. Conversely, BL specialists should predominate in blue open ocean waters at depth, where the light intensity is low, while CA4-capable strains would prevail in the upper mixed layer. The first hypothesis is indeed consistent with observations made along the *Tara* Oceans transect by Grébert and co-workers (26), since GL specialists were found to predominate at all depths in green waters in ocean margin areas. Yet these authors also reported a globally higher relative abundance of BL specialists in surface than at depth while the reverse was true for chromatic acclimaters. This global trend might however translate different local situations since, in temperate areas, surface and deep *Synechococcus* populations often belong to different lineages that may exhibit different PTs (11, 26).

This study also brought some novel insights concerning the perception of light quality by CA4-capable cells. Indeed, the variations of the PUB:PEB ratio observed over the course of the experiment in some light conditions, notably in LGL (Figs. S1 and S3), suggests that the CA4-capable strain perceived and quickly responded to a gradual change of the light quality occurring within the flask as the culture became denser, even though there was no change in ambient light color in the incubator. Interestingly, our results also revealed the previously unknown capacity of a CA4-capable strain to adjust its PE:PC ratio depending on light intensity (Fig. 4B), a behavior previously described in a CA3-capable strain but in response to change in light color (6). When grown in white light with either a red light or a GL specialist, the CA3-capable strain was indeed found to modify its PE:PC ratio to harvest the part of the light spectrum not used by its competitor, while keeping the total amount of these two pigments constant. In the present case, a possible explanation for the observed decrease of the PE:PC ratio of the CA4-capable strain with light intensity is that it may enable a modulation of energy transfer efficiency along the PBS rods.

## CONCLUSIONS

Previous studies predicted that flexible phenotypes would often be weaker competitors than specialists in mono-color conditions (Stomp *et al.*, 2008; Lovindeer *et al.*, 2021). Although we confirmed that specialists were the winners at LL in their favorite light color, and that the GL specialist was the best competitor in HGL, we also found that the CA4-capable strain was actually able to outcompete the other PTs in HBL.

Overall, our results demonstrated that both the light quality and quantity had major effects on the outcome of co-cultures experiments. Still, given the interplay between competition for the underwater light field and other important selective factors, such as nutrients or temperature, more studies are needed to better understand the spatio-temporal variability of the distribution of the different *Synechococcus* PTs at the global scale.

## AUTHOR CONTRIBUTIONS

**Louison Dufour:** Conceptualization; investigation; methodology; validation; visualization; formal analysis; data curation; writing – original draft. **Laurence Garczarek:** Conceptualization; funding acquisition; writing – review and editing; supervision; project administration. **Francesco Mattei:** Methodology; validation; writing – review and editing. **Bastian Gouriou:** Investigation; methodology; validation. **Julia Clairet:** Investigation; methodology; validation. **Morgane Ratin:** Investigation; methodology; validation. **David M. Kehoe:** Writing – review and editing. **Jef Huisman:** Methodology; validation; writing – review and editing. **Jolanda M. H. Verspagen:** Methodology; validation; writing – review and editing. **Frédéric Partensky:** Conceptualization; funding acquisition; formal analysis; writing – review and editing; supervision; project administration.

## ACKNOWLEDGEMENTS

We would like to thank members of the CRBM (Centre de Recherche en Biologie Marine, Roscoff) for sharing the temperature-controlled room and CO_2_ system used in this study, as well as Sarah Bureau for taking care of nutrient analyses. We are also most grateful to Christophe Six and Sarah Garric for technical hints on PAM fluorimetry, as well as Priscillia Gourvil, Martin Gachenot and Michele Grego from the Roscoff Culture Collection (http://roscoff-culture-collection.org/) for providing *Synechococcus* strains. This work was supported by the French “Agence Nationale de la Recherche” program EFFICACY (ANR-19-CE02-0019) granted to F.P. and L.G. and by United States National Science Foundation grants MCB-1029414 and MCB-1818187 to D.M.K. Figure 1 was created with Biorender.com.

## CONFLICT OF INTEREST STATEMENT

The authors have no conflicts of interest to declare.

---

**SUPPLEMENTARY FIGURE 1.**
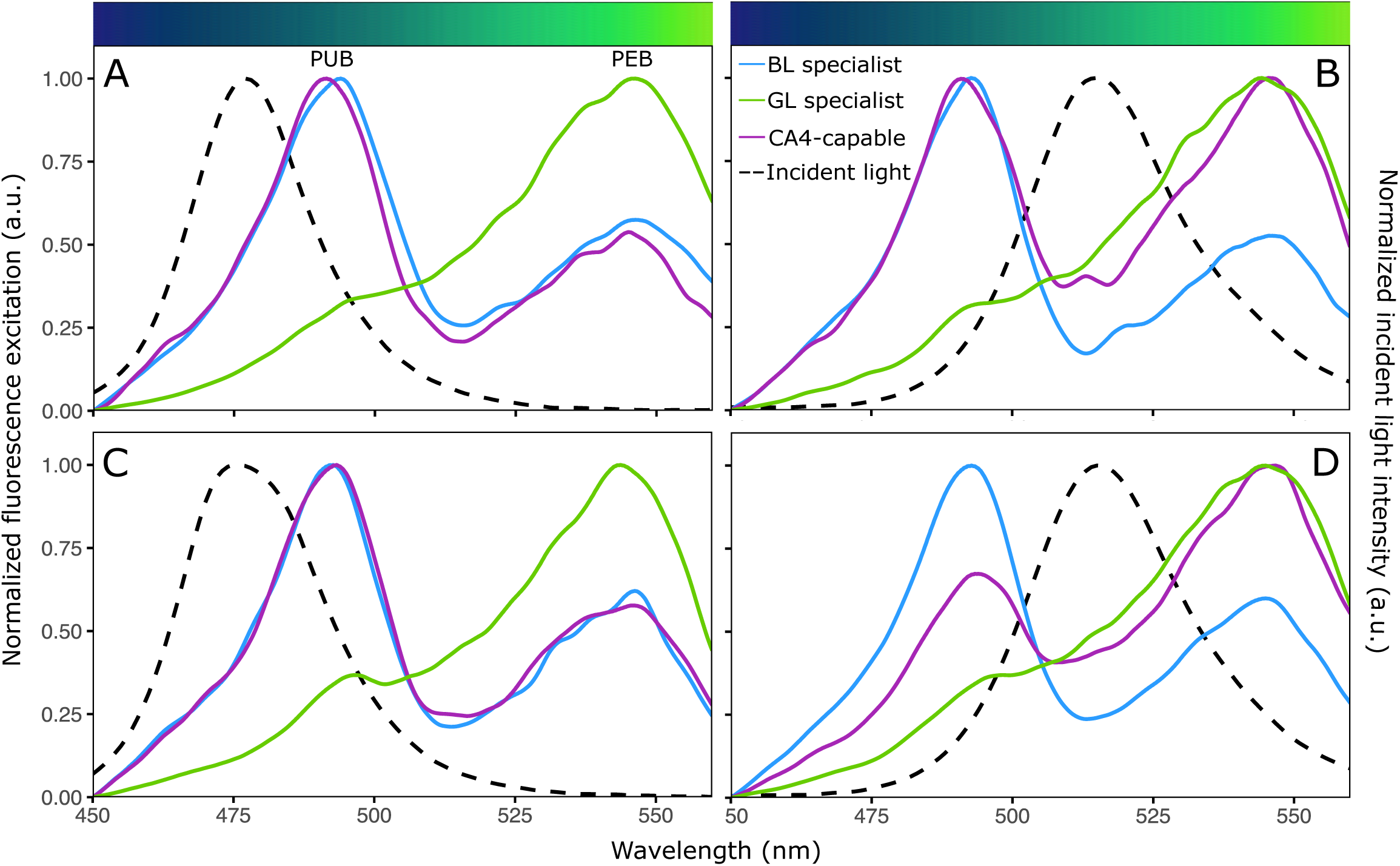
Representative excitation spectra of the three *Synechococcus* strains grown in the different light conditions used in this study. (A) Low blue light (LBL). (B) Low green light (LGL). (C) High blue light (HBL). (D) High green light (HGL). The LEDs spectra are represented by the black dashed line, while the excitation spectra of the green and blue specialists, as well as the CA4-capable strain, are illustrated in green, blue and purple solid lines, respectively.

**SUPPLEMENTARY FIGURE 2.**
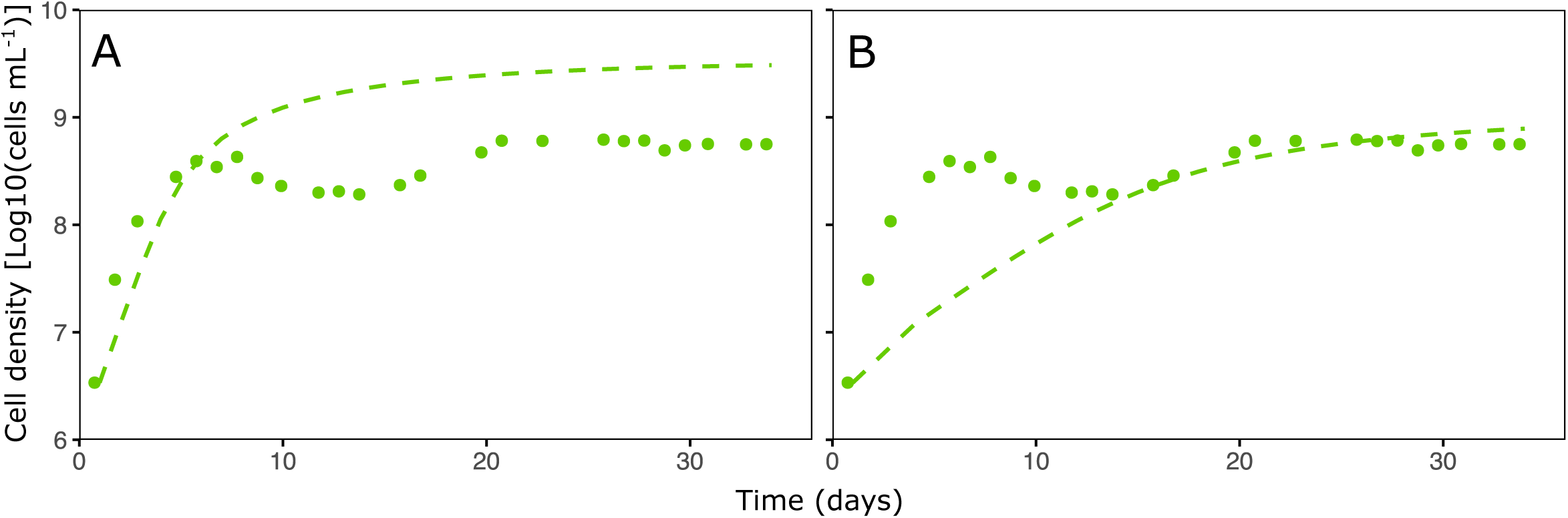
Example of model fitting to the data from the green light specialist in high green light (HGL). The model is able to capture either (A) the initial increase in population density but with an overestimation of the steady state population identity, or (B) the steady state population density but with an underestimation of the initial increase in population density. Points represent the data and lines represent the output of the model.

**SUPPLEMENTARY FIGURE 3.**
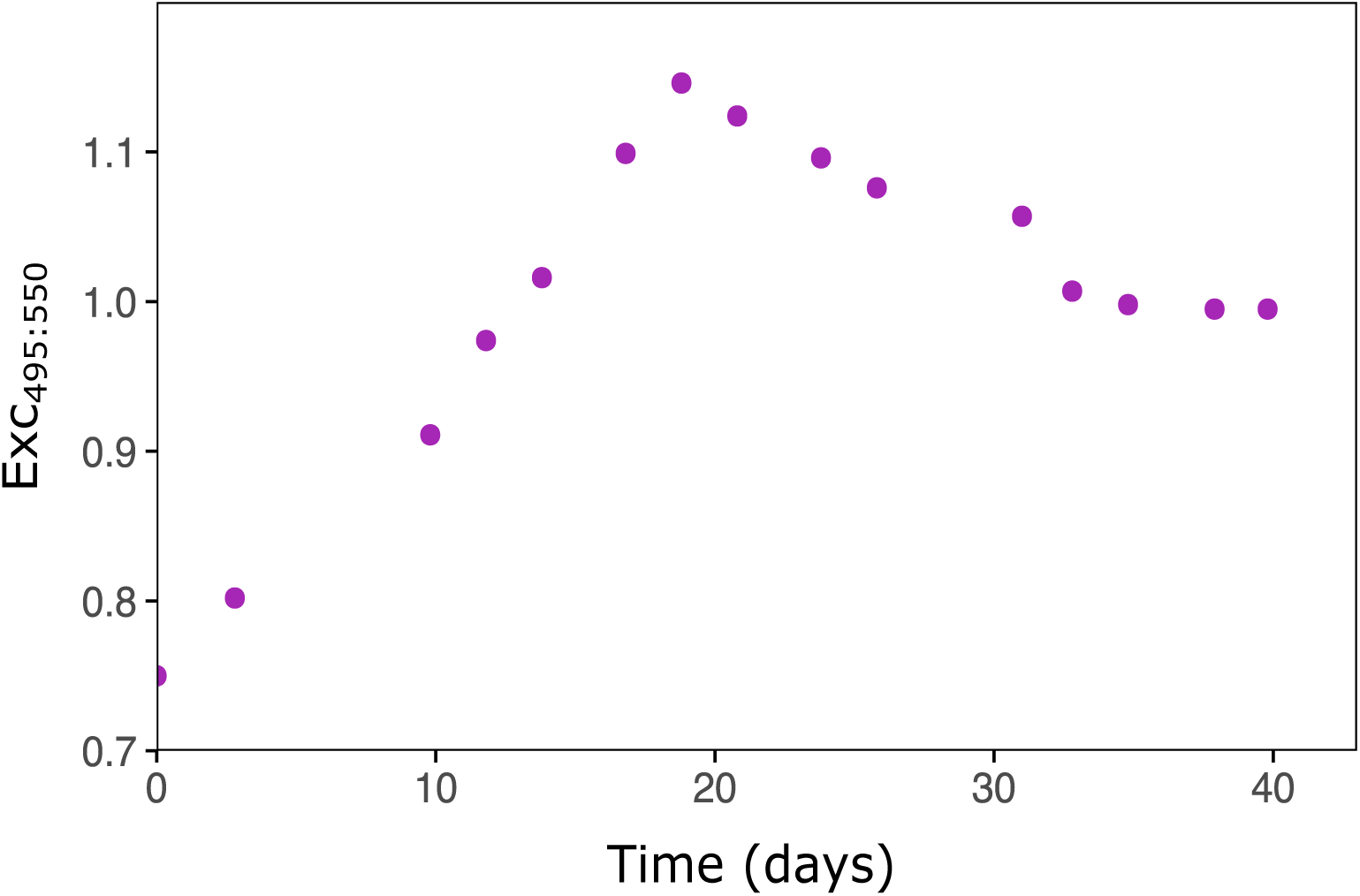
Time course variations of the Exc_495:550_ fluorescence excitation ratio, a proxy of the whole cell PUB:PEB ratio, for the CA4-capable strain in the low green light (LGL) monoculture.

**SUPPLEMENTARY FIGURE 4.**
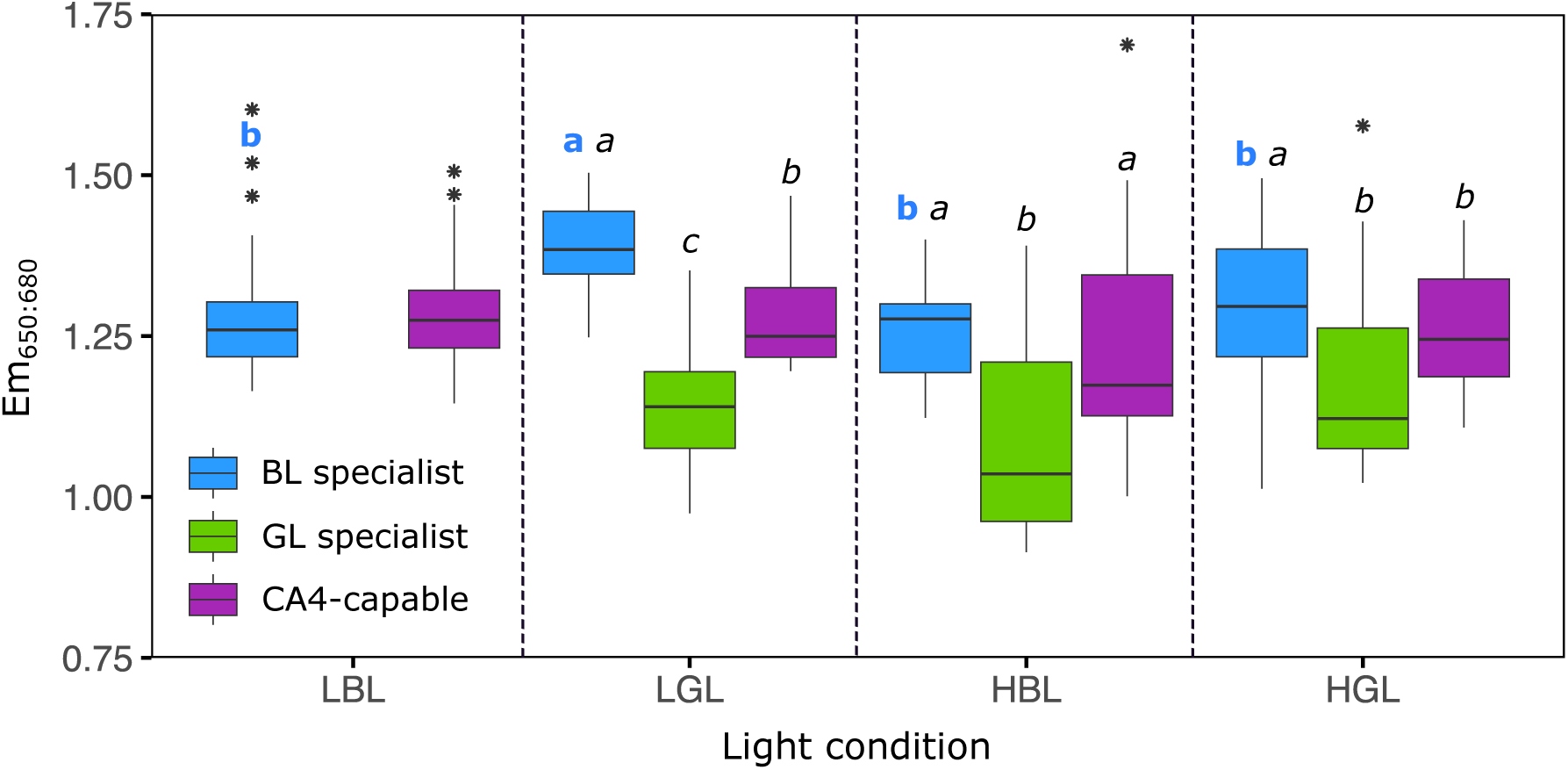
Mean Exc_650:680_ fluorescence emission ratio, a proxy of the whole cell phycocyanin to terminal acceptor (PC:TA) ratio, for the three *Synechococcus* strains grown in monoculture in the different light conditions used in this study. Boxplots represent all measurements performed over the course of the experiments. The different letters above boxplots indicate statistical test results (one-way ANOVA followed by Tukey’s test, or Kruskall-Wallis followed by Dunn’s test). The first bold colored letter compares the values displayed by a given strain in the four different light conditions tested. The second italic black letter compares the three different strains in a given light condition.

**SUPPLEMENTARY FIGURE 5.**
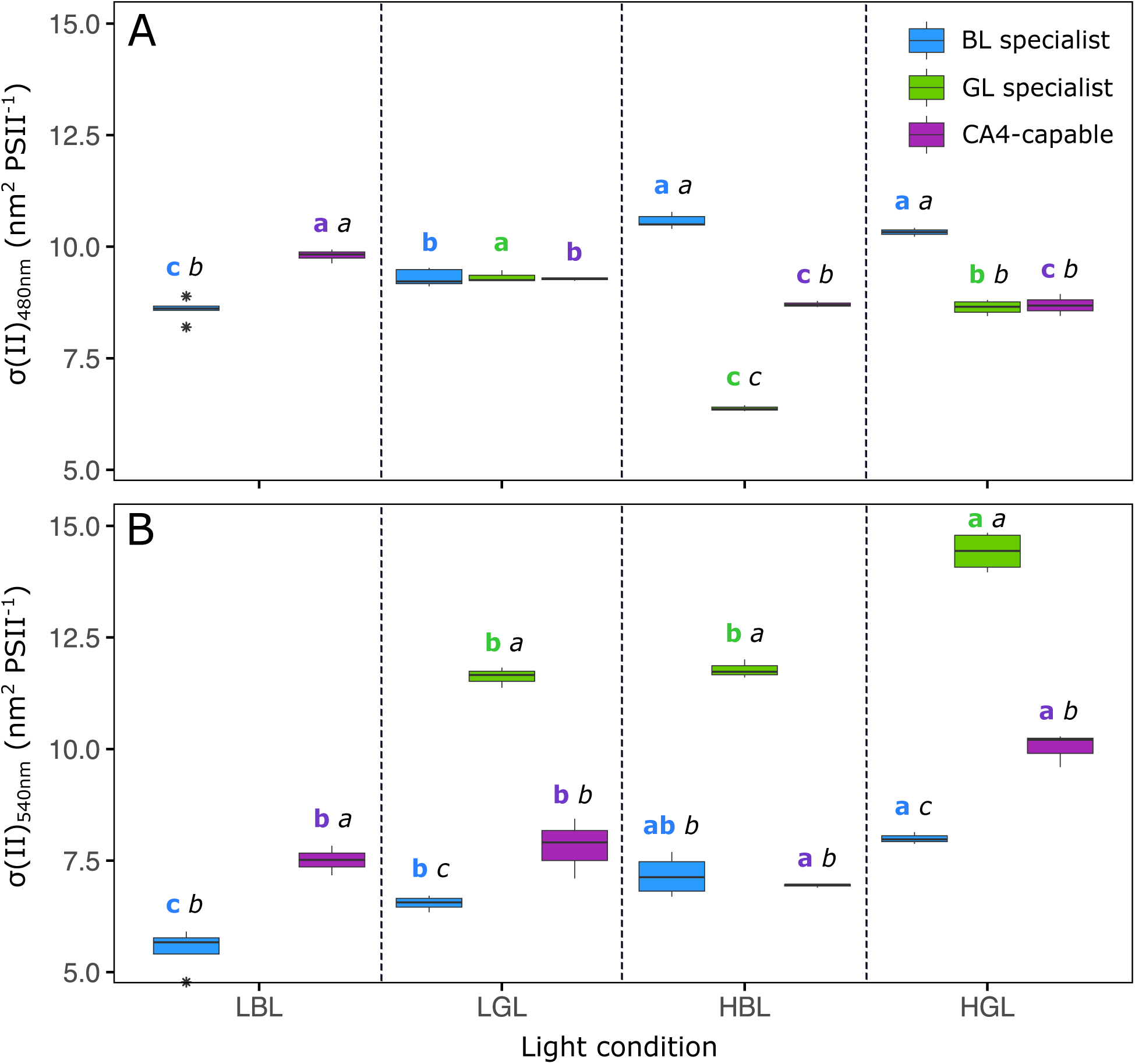
Photosystem II cross-section [σ(II)λ] for the three *Synechococcus* strains grown in monocultures in the different light conditions used in this study. (A) At 480 nm (cyan), the PAM excitation wavelength closest to PUB absorption peak (λ_max_ ≈ 495 nm). (B) At 540 nm (green), the PAM excitation wavelength closest to PEB absorption peak (λ_max_ ≈ 550 nm). Boxplots represent the ≥ 6 measurements performed during the growth phase and steady state for each strain and light condition. The different letters above boxplots indicate statistical test results for a specific wavelength (one-way ANOVA followed by Tukey’s test, or Kruskal-Wallis followed by Dunn’s test). The first bold colored letter compares the values displayed by a given strain in the four different light conditions tested. The second italic black letter compares the values exhibited by the three different strains in a given light condition.

**SUPPLEMENTARY FIGURE 6.**
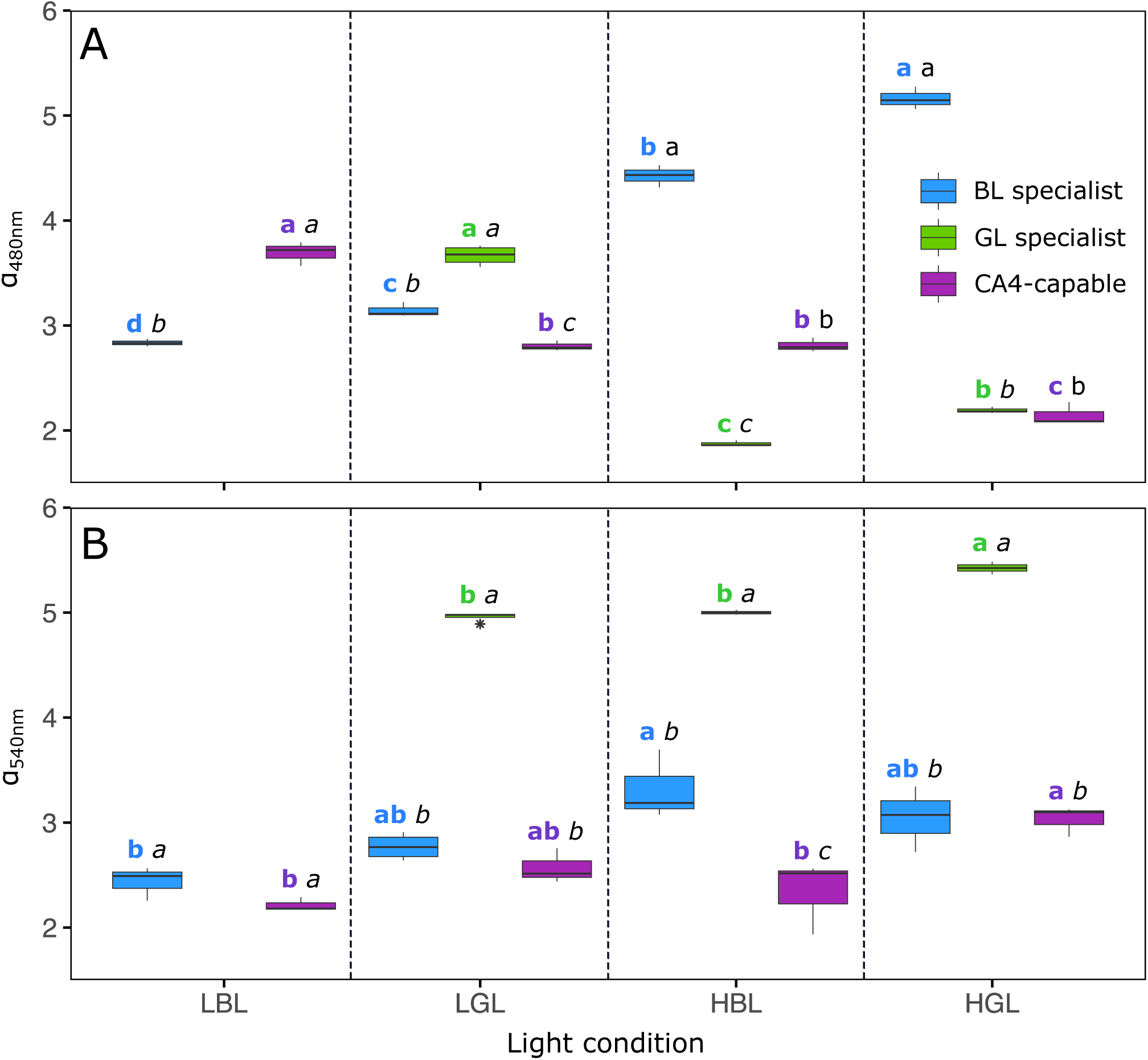
Same as Suppl. Fig. 5 but for photosystem II efficiency under non-saturating light (α).

